# Structure of the GOLD-domain seven-transmembrane helix protein family member TMEM87A

**DOI:** 10.1101/2022.06.20.496907

**Authors:** Christopher M. Hoel, Lin Zhang, Stephen G. Brohawn

## Abstract

TMEM87s are eukaryotic transmembrane proteins with two members (TMEM87A and TMEM87B) in humans. TMEM87s have proposed roles in protein transport to and from the Golgi, as mechanosensitive ion channels, and in developmental signaling. TMEM87 disruption has been implicated in cancers and developmental disorders. To better understand TMEM87 structure and function, we determined a cryo-EM structure of human TMEM87A in lipid nanodiscs. TMEM87A consists of a Golgi-dynamics (GOLD) domain atop a membrane spanning seven-transmembrane helix domain with a large cavity open to solution and the membrane outer leaflet. Structural and functional analyses suggest TMEM87A may not function as an ion channel or G-protein coupled receptor. We find TMEM87A shares its characteristic domain arrangement with seven other proteins in humans; three that had been identified as evolutionary related (TMEM87B, GPR107, and GPR108) and four previously unrecognized homologs (GPR180, TMEM145, TMEM181, and WLS)). Among these structurally related <u>GO</u>LD domain <u>s</u>even-<u>t</u>ransmembrane helix (GOST) proteins, WLS is best characterized as a membrane trafficking and secretion chaperone for lipidated Wnt signaling proteins. We find key structural determinants for WLS function are conserved in TMEM87A. We propose TMEM87A and structurally homologous GOST proteins could serve a common role in trafficking membrane-associated cargo.

## Introduction

TMEM87 proteins are transmembrane proteins found in eukaryotic organisms from fungi and plants to mammals^1^. In humans, two paralogs have been identified: TMEM87A and TMEM87B^1^. TMEM87A and TMEM87B have been associated with protein transport to and from the Golgi, mechanosensitive cation channel activity, and cardiac and other developmental processes^2–5^. In humans, TMEM87 proteins have been implicated in developmental diseases and cancers^6–9^.

Several lines of evidence support a role for TMEM87 in protein trafficking. First, TMEM87s localize to the Golgi^2,3^. Second, overexpression of TMEM87A or TMEM87B partially rescues defective endosome-to-*trans*-Golgi network retrograde traffic in HEK293 cells lacking the Golgi-associated retrograde protein (GARP) complex member VPS54^2^. Third, TMEM87A has been identified as retrograde cargo captured by mitochondrial golgins^3^. A recent study, however, associated TMEM87A with mechanosensitive cation channel activity^5^. Deflection of micropillar culture supports resulted in cationic currents in melanoma cells that were reduced by TMEM87A expression knockdown. Additionally, TMEM87A over-expression resulted in mechanosensitive currents in PIEZO1 knockout HEK293T cells^5^. TMEM87A knockout melanoma cells show increased adhesion strength and decreased migration, suggesting potential roles for TMEM87A in these processes in cancers^5^.

TMEM87B has been linked to developmental processes in recurrent 2q13 microdeletion syndrome^4,8,9^. Patients with this syndrome exhibit cardiac defects, craniofacial anomalies, and developmental delay. The vast majority of 2q13 microdeletions include TMEM87B^8^. In zebrafish, morpholino knockdown of TMEM87B resulted in cardiac hypoplasia^4^. In a patient with a severe cardiac phenotype, whole-exome sequencing uncovered a paternally inherited TMEM87B missense mutation and chromosome 2 deletion inherited from an unaffected mother^8^. Both TMEM87A and TMEM87B have additionally been implicated in cancers such as non-small cell lung cancer through fusion or suspected interaction with oncogenes^6,7,10^.

Bioinformatic analysis has grouped TMEM87A and TMEM87B in a small family of proteins (termed lung 7TM receptors or LUSTRs) with so-called orphan GPCRs GPR107 and GPR108^1^. Like TMEM87s, GPR107 and GPR108 localize to the Golgi and have suggested roles in protein trafficking. GPR107 is implicated in the transport of *P. aeruginosa* Exotoxin A, *C. jejuni* cytolethal distending toxin (CDT), and ricin, while GPR107 knockout cells display deficits in receptor mediated endocytosis and recycling^11–14^. GPR108 is critical for the transduction of a majority of AAV serotypes^15,16^. While a GPR107 knockout mouse was embryonic lethal, a GPR108 knockout mouse was viable and revealed a potential role for GPR108 in the regulation of Toll-like receptor (TLR) mediated signaling^14,17^.

Despite being implicated in important cell biological processes and human health, our molecular understanding of TMEM87s and LUSTR proteins is limited and no experimental structures of these proteins have been reported to date. Here, we present a cryo-electron microscopy (cryo-EM) structure of human TMEM87A. Through structural and bioinformatic analyses, we find TMEM87A belongs to a broader protein family we term <u>GO</u>LD-domain <u>s</u>even <u>t</u>ransmembrane (GOST) proteins. We speculate TMEM87A and other GOST proteins could function as trafficking chaperones for membrane associated cargo.

## Results

Full-length human TMEM87A was expressed and purified from Sf9 cells and reconstituted into lipid nanodiscs composed of MSP1D1 and a mixture of DOPE, POPS, and POPC lipids (Supplementary Fig. 1). Cryo-electron microscopy (cryo-EM) was used to determine the structure of TMEM87A to a nominal resolution of 4.7 Å, with better resolved regions reaching ~4.3 Å in the core of the protein. The map was of sufficient quality to place secondary structure elements unambiguously and, using an AlphaFold2 predicted structure as a starting model, to place and refine 395/555 residues (Fig. 1, Supplementary Fig. 2). The modeled portion of TMEM87A and corresponding predicted structure are similar (overall r.m.s.d. = 1.8 Å) with a minor difference in the relative position of extracellular and transmembrane regions. 37 N-terminal residues (amino acids 1-37), 88 C-terminal residues (475-555), and 36 residues in loops within the extracellular domain (145-173, 192-198) were not resolved in the cryo-EM map and are unmodeled.

**Figure 1.**
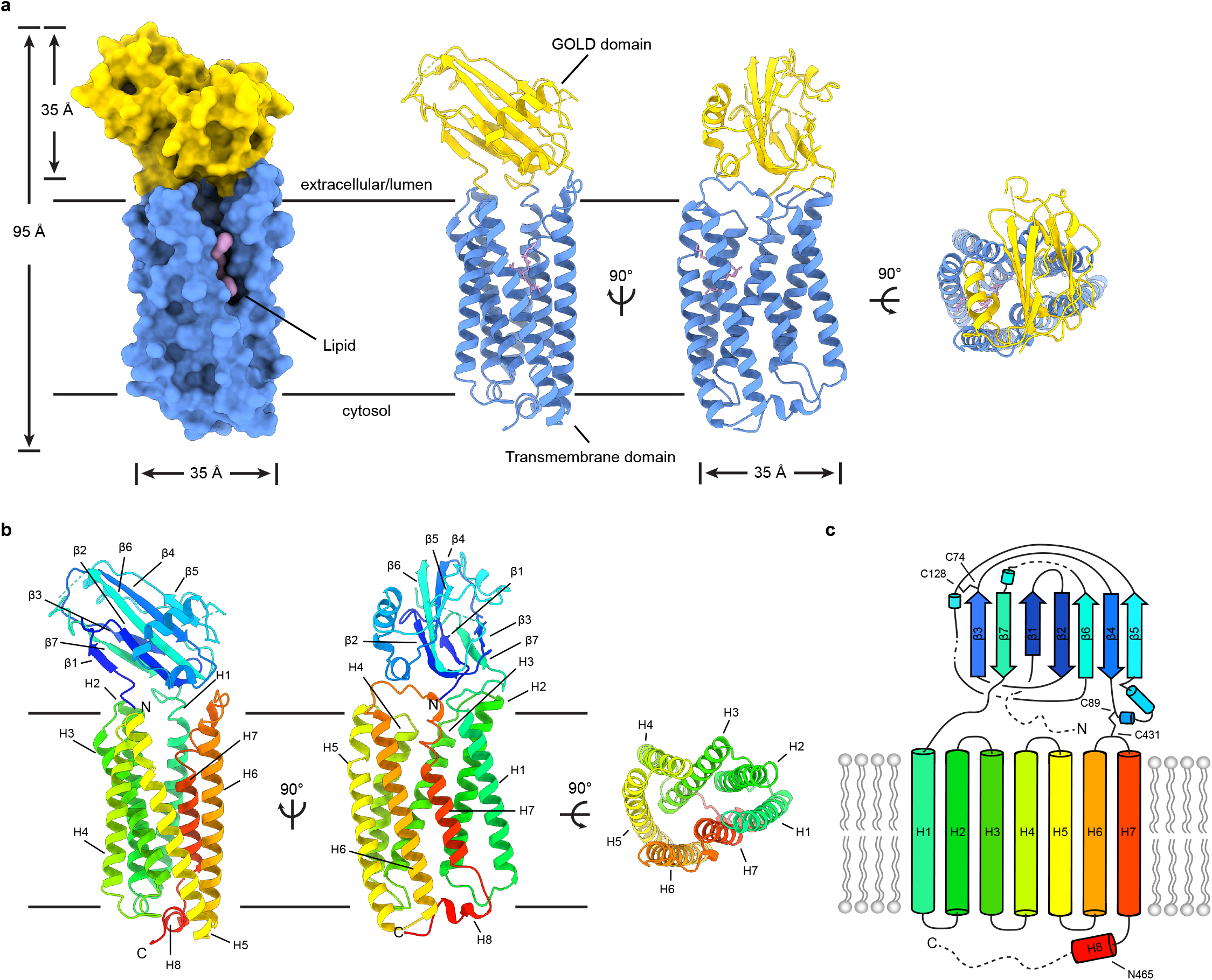
Structure of TMEM87A in lipid nanodiscs. **(a)** Model for TMEM87A viewed from the plane of the membrane (left) and the extracellular or lumenal side (right). The GOLD domain is colored yellow, seven-transmembrane domain colored blue, and phospholipid colored pink. **(b)** TMEM87A with rainbow coloring from N-terminus (blue) to C-terminus (red). **(c)** Corresponding cartoon of domain topology with rainbow coloring from N-terminus (blue) to C-terminus (red). Positions of disulfide bonds and residues noted in the text are indicated.

TMEM87A is composed of an extracellular beta-sandwich domain atop a G-protein coupled receptor (GPCR)-like seven transmembrane (7TM) domain (Fig. 1). The extracellular beta-sandwich domain is formed by seven beta strands in two opposing sheets, with three strands on the N-terminal face and four on the opposing face. The N-terminal face consists of β1, β3, β7, and a region between β5 and β6 lacking secondary structure that packs against β3 at the top of the sheet. The opposing face of the β-sandwich is formed from β2, β4, β5, and β6 with a helix-turn-helix motif between β4 and β5. A short 10 residue loop extends from the end of β7 to the 7TM domain at the extracellular side of H1. Two disulfide bonds are observed. The first, between C74 and C128, connects two loops at the top of this extracellular domain. The second, between C89 and C431, tethers the bottom of the extracellular domain to the top of the 7TM domain in the H6-H7 linker, perhaps constraining their relative movement.

The transmembrane region of TMEM87A is structurally similar to other 7TM proteins, with a few notable differences (Supplementary Fig. 3). Using Dali to compare the isolated TMEM87A transmembrane domain to all experimentally determined protein structures returns 7TM proteins as clear structural homologs, with the fungal class D GPCR Ste2, microbial opsins, and additional class A GPCRs among the hits^18–21^.

Since TMEM87A shares structural features with ion conducting opsins and previous work implicated TMEM87A in mechanosensitive cation conduction^5^, we asked whether TMEM87A displayed channel activity in isolation. We purified and reconstituted TMEM87A into proteoliposomes and recorded currents across patched membranes in response to membrane stretch induced by pressure steps. We observed neither basal nor mechanically activated currents in TMEM87A reconstituted proteoliposomes, in contrast to the mechanosensitive ion channel TRAAK used as a positive control (Supplementary Fig. 4). This result is consistent with the lack of a clear ion conducting path in the TMEM87A structure. We conclude that under these conditions, TMEM87A does not form a mechanosensitive ion channel.

Since TMEM87A contains a 7TM domain, we next asked whether it has structural features consistent with G-protein coupled receptors. TMEM87A and Class A GPCRs have a similar transmembrane helix organization, for example, TMEM87A and the β1-adrenergic receptor are superimposed with an overall r.m.s.d. of 4.13 Å (Fig. 2, Supplementary Fig. 3). Intriguingly, the position of cytoplasmic helix 8 in TMEM87A differs substantially from experimental structures of GPCRs. In TMEM87A, helix 8 turns back towards the center of the protein and packs against the bottom of the transmembrane domain (Fig. 2a,b). In experimental GPCR structures, helix 8 is instead rotated nearly 180° and adopts a different position projecting towards the surrounding membrane^22–24^ (Fig. 2b-c). This position of helix 8 in GPCRs appears necessary to accommodate G-protein or arrestin binding and is similarly positioned in apo- and complexed-receptor structures (Fig. 2c). The position of helix 8 in TMEM87A would sterically clash with putative G-protein or arrestin binding through interfaces analogous to those observed in GPCR structures (Fig. 2c-e). Notably, the TMEM87B point mutation implicated in 2q13 deletion syndrome (TMEM87B N456D) corresponds to TMEM87A residue N465 at the TM7-helix 8 junction^8^ (Fig. 2a), suggesting helix 8 is important for TMEM87B function. In addition to this difference in helix 8 position, TMEM87A does not share canonical motifs of class A or D GPCRs including the class A NPxxY activation motif, PIF motif, and D(E)/RY motif or the class D LPLSSMWA activation motif^19,25,26^. Taken together, these differences suggest TMEM87A may not couple to G proteins or arrestins like canonical GPCRs.

**Figure 2.**
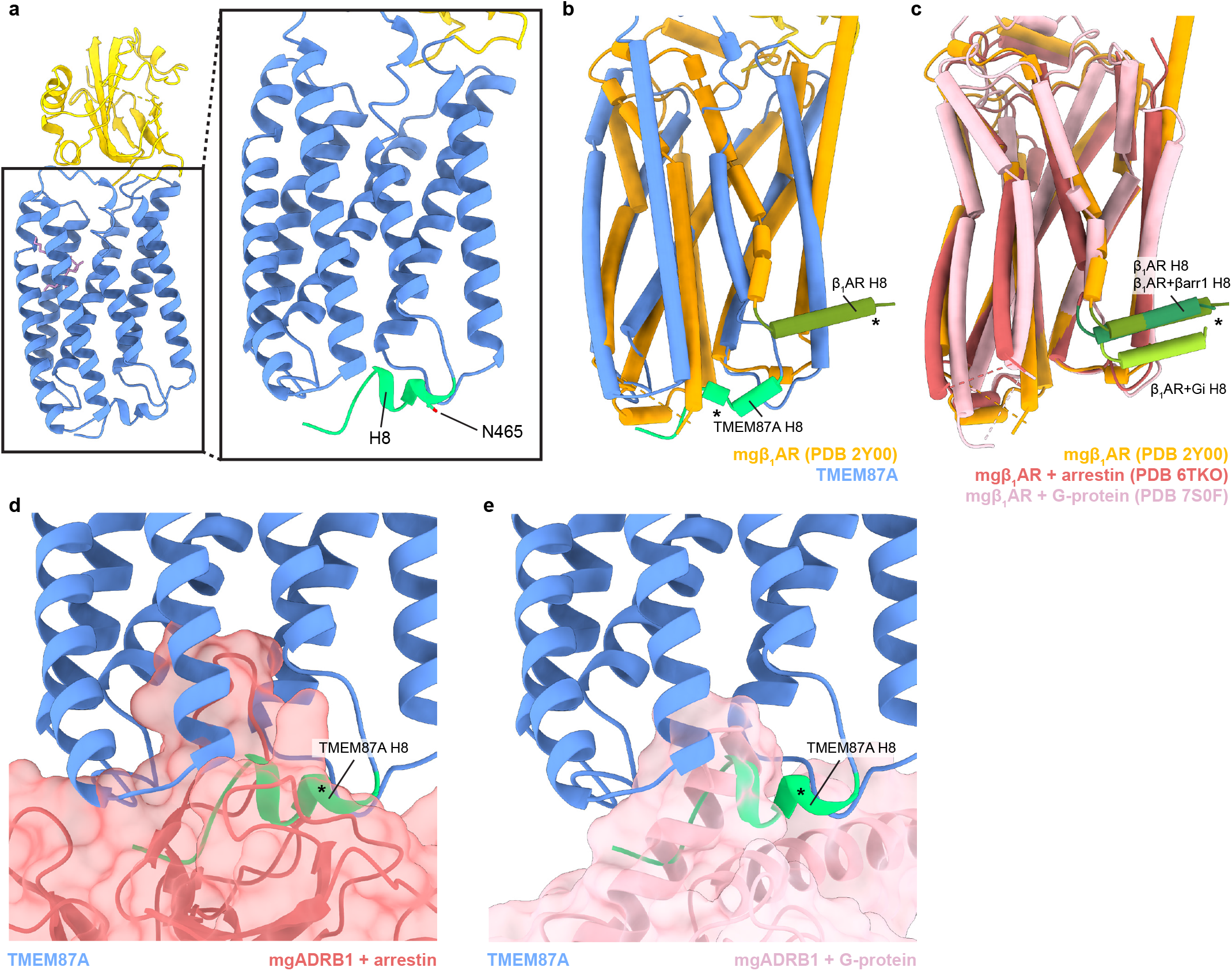
The TMEM87A transmembrane domain and comparison to an A class GPCR. **(a)** View of the TMEM87A model from the “back” in the plane of the membrane and zoomed view highlighting the position of helix 8 with residue N465 indicated (corresponding to TMEM87B N456D implicated 2q13 deletion syndrome). **(b)** Overlay of TMEM87A (blue) and β1 adrenergic receptor (mgβ1AR PDB: 2Y00, orange) transmembrane domains. **(c)** Overlay of mgβ1AR alone (PDB: 2Y00, orange), in complex with arrestin (PDB: 6TKO, red) in complex with heterotrimeric Gi-proteins (PDB: 7S0F, pink). **(d,e)** Overlay of TMEM87A (blue) and **(d)** arrestin (red surface) or **(e)** heterotrimeric Gi (pink surface) from mgβ1AR complex structures(PDB: 6TKO, 7S0F). Helix 8s (H8) are colored shades of green and denoted with asterisks.

We found the extracellular/lumenal region of TMEM87A adopts a Golgi dynamics (GOLD) domain fold. Using Dali to compare the isolated TMEM87A extracellular β-sandwich domain (residues 38-218) to all experimentally determined protein structures returns human p24 GOLD domain proteins as top hits. Superposition of the TMEM87A extracellular domain and p24 shows conservation of β strand topology throughout the domain, with differences predominantly in regions connecting β strands including the helixturn-helix motif in the TMEM87A GOLD domain^18,27^ (Supplementary Fig. 5a-c). GOLD domain proteins have established roles in the secretory pathway^27,28^ and p24 proteins are implicated as cargo receptors for protein transport, with the GOLD domain mediating cargo recognition^29,30^. Among different p24 proteins, the GOLD domain loops are among the most variable regions and have been hypothesized to recognize different cargo^27^. Identification of a GOLD domain in TMEM87A suggests a structural explanation for proposed roles of TMEM87s in protein transport, with major differences in GOLD domain loops between TMEM87A and p24 perhaps indicative of differences in interacting partners.

We next asked whether further structural comparison could provide insight into TMEM87A functions. We performed a Dali search against the entire AlphaFold2-predicted human proteome using the experimental TMEM87A structure as the reference^18,31,32^. This search uncovered proteins predicted to have the same domain topology consisting of a GOLD domain fused to a 7TM domain: all four members of the LUSTR family and four previously unrecognized homologs GPR180, TMEM145, TMEM181, and Wntless (WLS) (Fig. 3, Supplementary Fig. 5d-j, Supplementary Fig. 6a-g). The predicted structures are all structurally homologous to TMEM87A (overall r.m.s.d range from 1.85 to 3.87 Å). We propose an expansion of the LUSTR family to include these additional members and to rename them GOST proteins (for <u>GO</u>LD domain <u>s</u>even <u>t</u>ransmembrane helix proteins).

**Figure 3.**
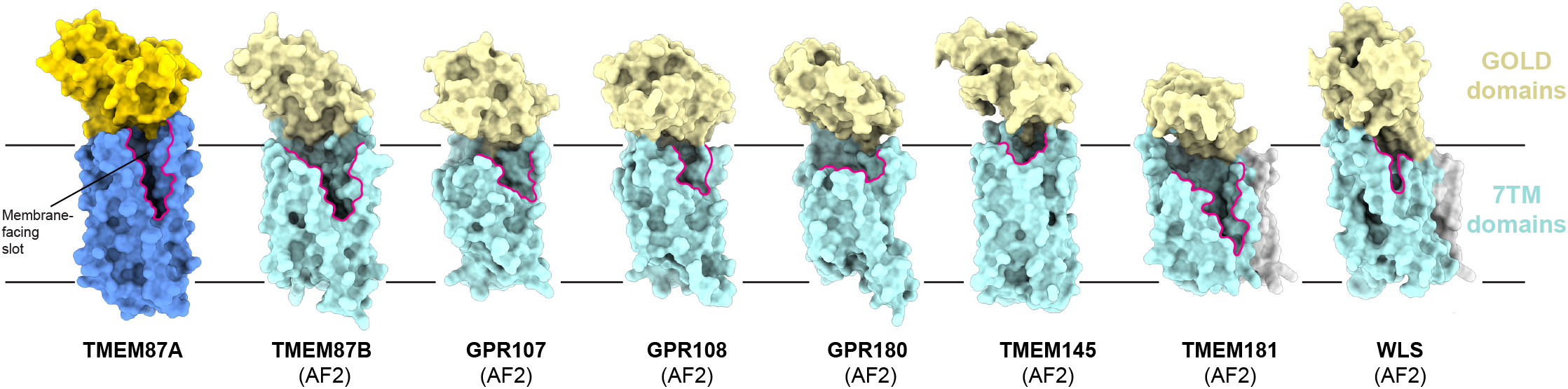
Predicted structural homology of GOST (GOLD domain seven transmembrane helix) family proteins. Experimentally determined TMEM87A structure and AlphaFold2 predicted structures of all identified human GOST family proteins (TMEM87B, GPR107, GPR108, GPR180, TMEM145, TMEM181, and WLS) shown as surfaces in the same orientation from the membrane plane. GOLD domains are yellow, seven-transmembrane domains are blue, N-terminal helices (if present) are gray, and membrane-facing slots are outlined in pink. Low confidence (<80 pLDDT) regions of predicted structures are removed for clarity (full structures are displayed in Supplementary Fig. 6).

While three newly identified GOST proteins have been minimally characterized to date (prior work has associated GPR180 with TGF-B signaling and TMEM181 with *E. coli* cytolethal distending toxin toxicity^33,34^), WLS (also known as GPR177 or evenness interrupted homolog (EVI)) has been extensively studied. WLS plays an essential role in chaperoning lipidated and secreted Wnt signaling proteins between internal membranes and the cell surface^35–37^. Recent cryo-EM structures of WLS bound to Wnt3a or Wnt8a provide insight into the basis for chaperone-client interactions^38,39^ (Fig. 4a). The extracellular region of Wnt interacts extensively with the WLS GOLD domain and an extended palmitoleated hairpin of Wnt that is buried in the large hydrophobic cavity within the WLS 7TM domain. Comparison of TMEM87A with WLS shows that the transmembrane regions are well aligned with two notable differencs. First, WLS H4 and H5 are shifted approximately 13 Å away from the center of the protein, resulting in a larger cavity compared to TMEM87A necessary to accommodate the insertion of the lipidated Wnt hairpin (Fig. 4b). The relative orientation of the GOLD and 7TM domains in TMEM87A and WLS also differs, with the WLS GOLD domain shifted approximately 26 Å in the same direction of movement as the H4/H5 shift (Fig. 4b). Intriguingly, TMEM87A and WLS exhibit similar overall patterns of surface electrostatic charge including in the 7TM cavity that, in WLS, mediates binding of the Wnt lipidated hairpin (Fig. 4c-d).

**Figure 4.**
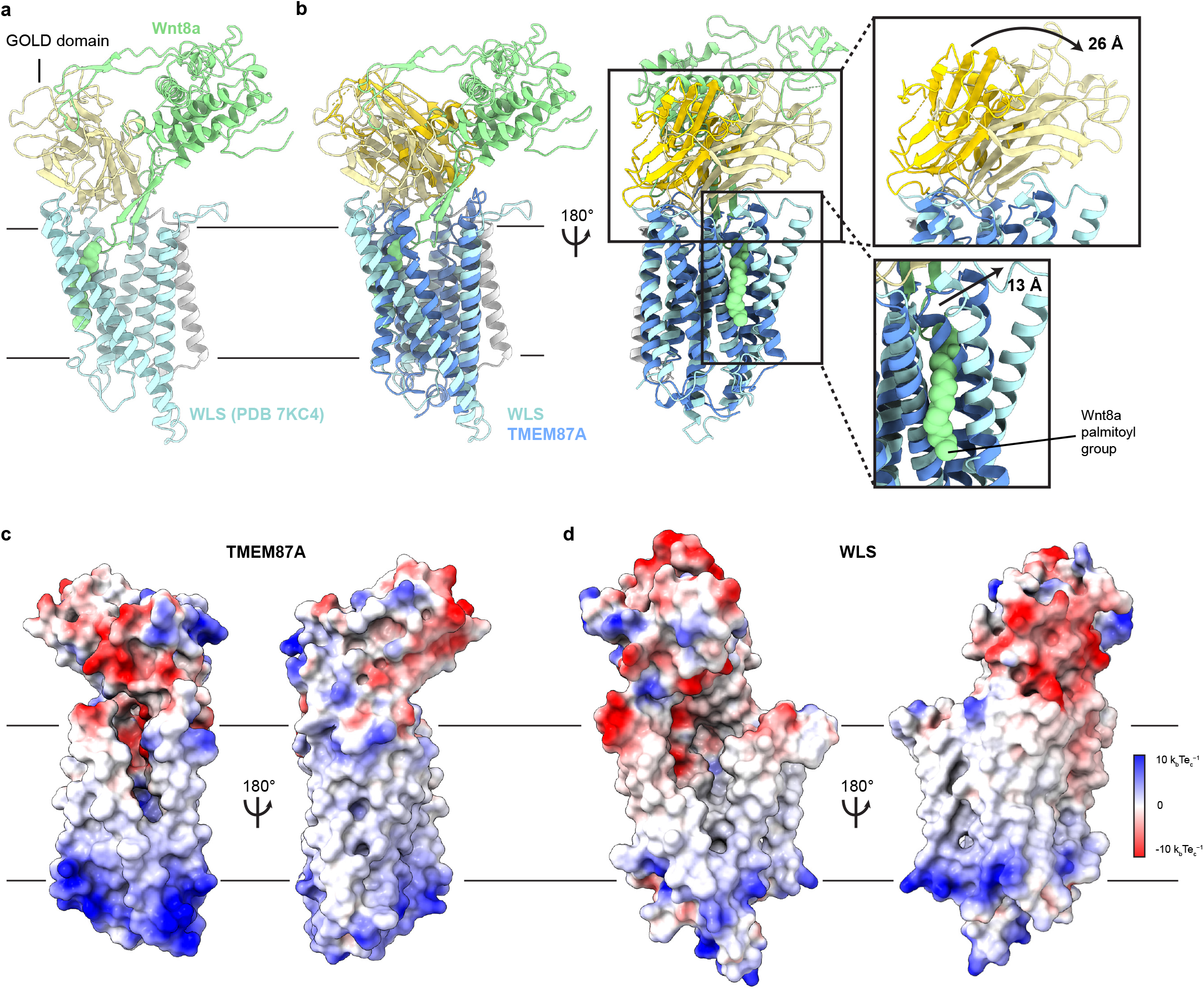
Structural comparison of TMEM87A and WLS bound to Wnt. **(a)** WLS-Wnt8a complex structure (PDB: 7KC4) with WLS GOLD domain light yellow, the WLS seven-transmembrane domain light blue, the N terminal helix light gray, and Wnt8a light green. **(b)** Overlay of TMEM87A and WLS (PDB 7KC4) experimental structures, with TMEM87A GOLD domain colored yellow and TMEM87A transmembrane domain colored blue. Zoomed in views highlight the differences in relative domain orientation between TMEM87A and WLS. **(c,d)** Views of **(c)** TMEM87A and **(d)** WLS electrostatic surfaces. Scale is from −10 k_b_Te_c_^−1^ to 10 k_b_Te_c_^−1^.

Perhaps the most striking structural feature of TMEM87A is the large cavity within the 7TM domain, a feature that is similar in several respects to the cavity in WLS structures (Fig. 5a,d). In TMEM87A this cavity measures approximately 8637 Å^3^ as compared to 12,350 Å^3^ for the WLS cavity. It is funnel shaped, presenting a large opening to the extracellular side near the GOLD domain and tapering closed near the intracellular surface above helix 8. The corresponding cavity in experimental WLS structures is filled by the lipidated Wnt3a/8a hairpin (Fig. 5d). Interestingly, the predicted WLS structure shows a smaller cavity relative to experimental WLS-Wnt structures. This suggests the apo-WLS cavity could dilate to accommodate Wnt binding (Supplementary Fig. 6h-j). The TMEM87A cavity is also exposed to the lipid membrane. H5 and H6 are splayed apart to open a large “slot” between the cavity and the upper leaflet of the bilayer (Fig. 5b). Density within the membrane exposed portion of the cavity is consistent with a bound phospholipid. Notably, the WLS cavity is open to the upper leaflet through a similar slot between H5 and H6 and a phospholipid is bound in a similar position in experimental structures (Fig. 5d-f). The slot in WLS is presumably important because it permits access of the lipidated Wnt hairpin from the membrane into the cavity binding site. Mapping sequence conservation onto TMEM87A and WLS structures shows a similar pattern with strongly conserved residues in the transmembrane region lining the internal cavity (Supplementary Fig. 7). Taken together, the structural homology between TMEM87A and WLS suggests they could similarly have evolved to bind and facilitate membrane trafficking of hydrophobic cargo.

**Figure 5.**
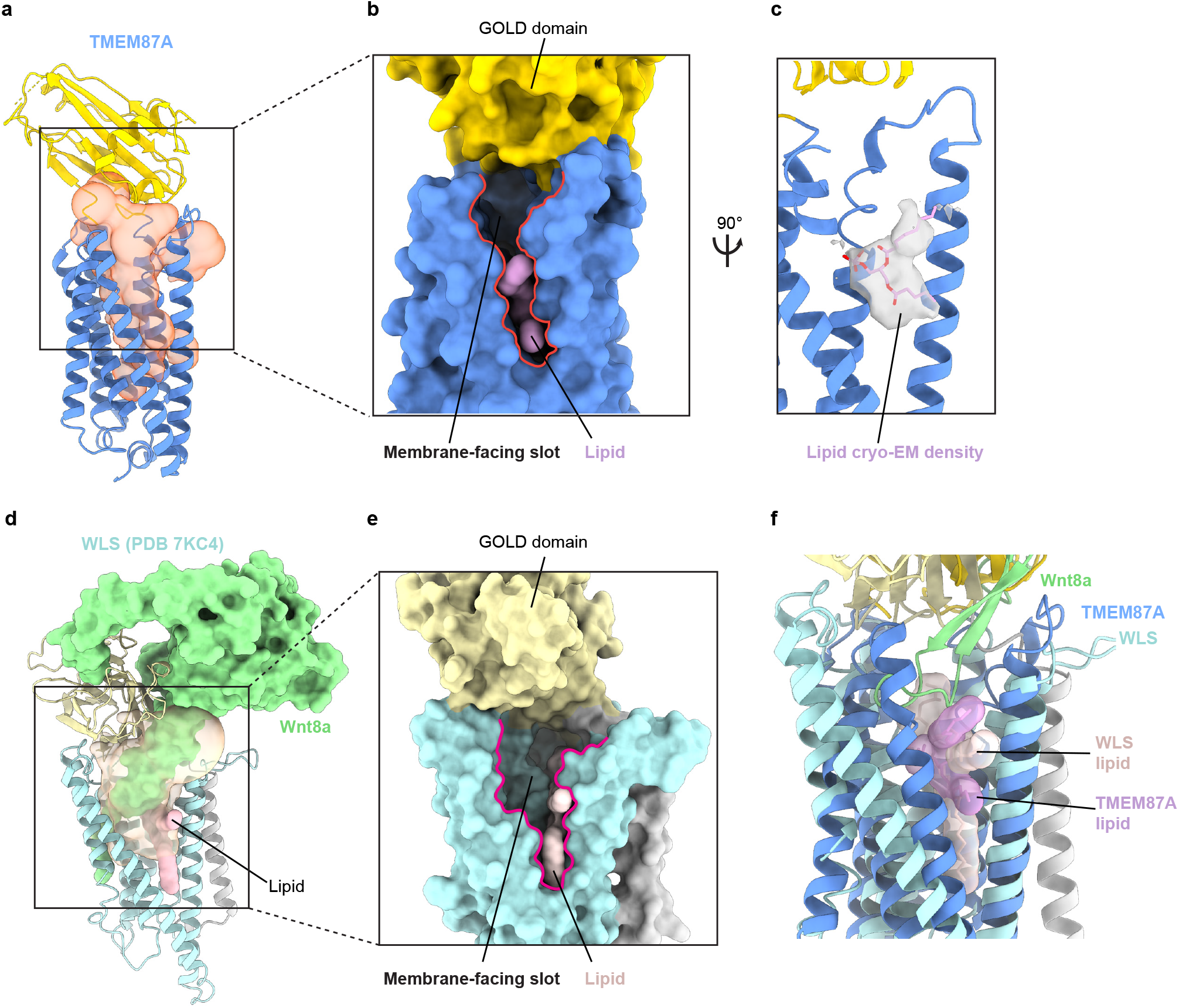
TMEM87A and WLS cavities are exposed to solution and membrane. **(a)** TMEM87A structure with CASTp calculated internal cavity shown as a transparent orange surface. **(b)** Surface of TMEM87A with membrane-facing slot (red outline) connecting the cavity and upper leaflet of the bilayer highlighted. **(c)** Modeled phospholipid (pink) is shown as a surface (left) or within cryo-EM density (right). **(d)** WLS (PDB: 7KC4) structure with CASTp calculated internal cavity shown as a transparent yellow surface, with bound Wnt8a (pale green surface) and phospholipid (pink) shown filling the cavity. **(e)** Zoomed in view shows surface representation of WLS (PDB: 7KC4) with open membrane-facing slot (red outline) and phospholipid (pink). **(f)** Overlay of the experimental TMEM87A and WLS structures showing relative positions of the H5-H6 membrane-facing slot and modeled phospholipids.

## Discussion

In this study, we report the first experimental structure of TMEM87A and consider its potential functional roles. TMEM87A shows general structural homology to ion transporting or conducting opsins and a recent report proposed TMEM87A contributes to mechanosensitive ion channel activity in cultured cells^5^. However, we found no evidence of channel activity in reconstituted proteoliposomes. This is consistent with the structure because while TMEM87A contains a solvent exposed cavity in its 7TM domain, it is sealed to the cytoplasmic side and open to the lipid bilayer, unlike typical ion conduction pathways. We cannot exclude the possibility that TMEM87A contributes to mechanosensitive ion channel currents under other conditions or in complex with other proteins. Still, further analyses suggest it may be more likely to act indirectly, perhaps through regulating transport, expression, or activity of a mechanosensitive ion channel.

We discovered that TMEM87 consists of an extracellular or lumenal <u>GO</u>LD domain atop a membrane spanning <u>s</u>even <u>t</u>ransmembrane helix domain and shares this characteristic organization with at least seven other GOST proteins in humans. Similarity to canonical GPCRs in the transmembrane region raises the question of whether TMEM87A or other GOST proteins are competent for GPCR signaling. While ligands have been proposed for some GOST proteins, specifically, neurostatin for GPR107 and L-lactate for GPR180, no definitive molecular evidence of G-protein signaling has been reported for any family member and GPR180 has been reported not to signal as a GPCR^33,40,41^. The position of helix 8 in TMEM87A is sterically incompatible with G-protein or arrestin binding and predicted structures of other GOST proteins present similar steric blocks. Together, this raises the possibility that GOST proteins are structurally incompatible with canonical GPCR signaling.

If GOST proteins do not function as channels or GPCRs, what functional roles might they serve? Most GOST family proteins are relatively understudied, though previous work supports roles for six members in different aspects of protein transport^2,3,11–16,34–36^. The best studied is WLS, for which a large body of work describes an essential role in Wnt signaling. Wnts must be secreted from the cell to accomplish their signaling functions, but they are lipidated during maturation and therefore embedded within intracellular membranes. WLS facilitates Wnt transport and secretion by interacting simultaneously with hydrophilic Wnt extracellular regions (through its GOLD domain) and hydrophobic Wnt membrane-embedded regions (though the 7TM cavity) Structurally, TMEM87A and other GOST proteins display, or are predicted to display, key features suggesting they could serve analogous roles; they (i) have GOLD domains positioned to interact with extracellular soluble domains or client proteins, (ii) have large transmembrane cavities, and (iii) expose the cavities simultaneously to the extracellular side and the membrane. This raises the intriguing possibility that GOST family proteins could transport lipidated or otherwise hydrophobic secreted proteins through a mechanism analogous to Wnt binding and transport by WLS. In this scenario, the two domains of GOST proteins would perform a sort of “coincidence detection” in which the GOLD domain binds solvent-exposed portions and the internal cavity binds membrane-buried portions of cargo molecules. Potential cargo for GOST proteins includes Wnts and other lipidated secreted proteins such as ghrelin and some cytokines^42–44^. Future studies are necessary to test the hypothesis that the GOST family proteins have common roles in transport of lipidated or membrane embedded proteins.

## Methods

### Cloning, expression and purification

The coding sequence for *Homo sapiens* TMEM87A (Uniprot ID: Q8NBN3) was codon-optimized for expression in *Spodoptera frugiperda* cells (Sf9 cells) and synthesized (Integrated DNA Technologies). The sequence was cloned into a custom vector based on the pACEBAC1 backbone (MultiBac, Geneva Biotech) with an added C-terminal PreScission protease (PPX) cleavage site, linker sequence, superfolder GFP (sfGFP) and 7×His tag, generating a construct for expression of TMEM87A-SNS-LEVLFQGP-SRGGSGAAAGSGSGS-sfGFP-GSS-7×His. MultiBac cells were used to generate a Bacmid according to the manufacturer’s instructions. Sf9 cells were cultured in ESF 921 medium (Expression Systems) and P1 virus was generated from cells transfected with FuGENE transfection reagent (Active Motif) according to the manufacturer’s instructions. P2 virus was then generated by infecting cells at 2 million cells per ml with P1 virus at a multiplicity of infection of roughly 0.1, with infection monitored by fluorescence and harvested at 72 h. P3 virus was generated in a similar manner to expand the viral stock. The P3 viral stock was then used to infect Sf9 cells at 2 million cells per ml at a multiplicity of infection of around 2~5. At 72 h, infected cells containing expressed TMEM87A-sfGFP protein were collected by centrifugation at 1,000*g* for 10 min and frozen at −80 °C. A cell pellet from 1 L culture was thawed and lysed by sonication in 100 mL buffer containing 50 mM Tris pH 8.0, 150 mM NaCl, 1 mM EDTA and protease inhibitors (1 mM phenylmethysulfonyl fluoride, 1 μM E64, 1 μg/mL pepstatin A, 10 μg/mL soy trypsin inhibitor, 1 μM benzamidine, 1 μg/mL aprotinin, and 1 μg/mL leupeptin). The membrane fraction was collected by centrifugation at 150, 000g for 45 min and homogenized with a cold Dounce homogenizer in 100 mL buffer containing 20 mM Tris pH 8.0, 150 mM NaCl, 1 mM EDTA, 1% n-dodecyl-b-D-maltopyranoside (DDM), 0.2% cholesteryl hemisuccinate (CHS) and protease inhibitors. Protein was extracted with gentle stirring for 2 h at 4 °C. The extraction mixture was centrifuged at 33,000g for 45 min and the supernatant was bound to 5 mL Sepharose resin coupled to anti-GFP nanobody for 2 h at 4 °C. The resin was collected in a column and washed with 25 mL buffer 1 (20 mM Tris pH 8.0, 150 mM NaCl, 1 mM EDTA, 0.025% DDM, 0.005% CHS), 50 mL buffer 2 (20 mM Tris pH 8.0, 500 mM NaCl, 1 mM EDTA, 0.025% DDM, 0.005% CHS) and 25 mL buffer 1. The resin was then resuspended in 5 mL of buffer 1 with 0.5 mg PPX protease and rocked gently in the capped column overnight. Cleaved TMEM87A was eluted with an additional 8 mL buffer 1, spin concentrated to roughly 500 μL with Amicon Ultra spin concentrator 50-kDa cutoff (Millipore), and then loaded onto a Superose 6 increase column (GE Healthcare) on an NGC system (Bio-Rad) equilibrated in buffer 1. Peak fractions containing TMEM87A were then collected and spin concentrated before incorporation into proteoliposomes or nanodiscs.

### Nanodisc reconstitution

Freshly purified TMEM87A was reconstituted into MSP1D1 nanodiscs with a mixture of lipids (DOPE:POPS:POPC at a 2:1:1 mass ratio, Avanti) at a final molar ratio of 1:4:400 (TMEM87A:MSP1D1:lipid mixture)^45^. First, 20 mM solubilized lipids in nanodisc formation buffer (20 mM Tris pH 8.0, 150 mM NaCl, 1 mM EDTA) was mixed with additional DDM detergent and TMEM87A. This solution was mixed at 4 °C for 30 min before addition of purified MSP1D1. The solution with MSP1D1 was mixed at 4 °C for 30 min before addition of 200 mg of Biobeads SM2. This mix was incubated at 4 °C for 30 min before addition of another 200 mg of Biobeads. This final mixture was then gently tumbled at 4 °C overnight (roughly 12 h). Supernatant was cleared of beads by letting large beads settle and carefully removing liquid with a pipette. Sample was spun for 10 min at 21,000g before loading onto a Superose 6 increase column in buffer containing 20 mM Tris pH 8.0, 150 mM NaCl. Peak fractions corresponding to TMEM87A in MSP1D1 were collected, 50-kDa cutoff spin concentrated and used for grid preparation. MSP1D1 was prepared as described without cleavage of the His-tag.

### EM sample preparation and data collection

TMEM87A in MSP1D1 nanodiscs was centrifuged at 21,000 × g for 5 min at 4 °C. A 3 μL sample was applied to holey carbon, 300 mesh R1.2/1.3 gold grids (Quantifoil, Großlöbichau, Germany) that were freshly glow discharged for 25 s. Sample was incubated for 5 s at 4 °C and 100% humidity prior to blotting with Whatman #1 filter paper for 3 s at blot force 1 and plunge-freezing in liquid ethane cooled by liquid nitrogen using a FEI Mark IV Vitrobot (FEI/Thermo Scientific, USA). Grids were clipped and transferred to a FEI Talos Arctica electron microscope operated at 200 kV. Fifty frame movies were recorded on a Gatan K3 Summit direct electron detector in super-resolution counting mode with pixel size of 0.5575 Å. The electron dose rate was 8.849 e^−^ Å^2^ s^−1^ and the total dose was 50.0 e^−^ Å^2^. Nine movies were collected around a central hole position with image shift and defocus was varied from −0.6 to −1.8 μm through SerialEM^46^.

### Cryo-EM data processing

Motion-correction with dose-weighting was performed using RELION3.1’s implementation of MotionCor2 and “binned” 2x from super-resolution to the physical pixel size^47–49^. CTFFIND-4.1 was then used to estimate the contrast transfer function (CTF)^50^. Micrographs with a CTF maximum estimated resolution lower than 5 Å were discarded. Particle images were picked first with RELION3.1’s Laplacian-of-Gaussian filter, then following initial clean-up and 2D-classification, templated-based auto-picking was performed. For this particle set, 2Dclassification was iteratively performed in both RELION3.1 and cryoSPARC v2, then iterative ab initio and heterogeneous refinement jobs were used to further identify higher quality particles^51^. Once an initial high quality set of particles was determined, these particle positions were used for training in Topaz, and the resulting Topaz model used to repick particles^52^. The above pipeline was applied again to the Topaz picked particles and the resulting set of particles was merged with the initial set of template-picked particles, and duplicates removed. This final set of particles was then input to Bayesian particle polishing in RELION3.1. The output “shiny” particles were then input to iterative homogenous and nonuniform refinements in cryoSPARC v2 until no further improvements were observed^53^. The output of the best nonuniform refinement was used for particle subtraction in RELION3.1, after which final refinement and postprocessing jobs were performed. This final refinement was input to Phenix resolve density modification to generate an additional map used during modeling^54^. The initial resolution and dynamic mask nonuniform parameters were adjusted empirically to yield the best performing refinement. UCSF pyem was used for conversion of files from cryoSPARC to Relion formats^64^.

### Model building and refinement

The final relion postprocessed map and Phenix density modified map were used for modeling. The AlphaFold2 TMEM87A model was rigid body fit to the density in ChimeraX and used as a foundation for manual model building in Coot^55,56^. The model was real space refined in Phenix and assessed for proper stereochemistry and geometry using Molprobity^57,58^. FSCs calculated in Phenix mtriage. Structures were analyzed and figures were prepared with CASTp, DALI, ChimeraX, JalView, Prism 8, Python GNU Image Manipulation Program, and Adobe Photoshop and Illustrator software^18,59,60^. Consurf was used to map conservation onto the structure surfaces using an alignment of sequences determined using SHOOT^61,62^. Electrostatic potentials calculated in ChimeraX. CASTp cavity calculation output was converted from JSON to PDB using a custom python script.

### Proteoliposome reconstitution

For proteoliposome patching experiments, we incorporated protein into lipids and generated proteoliposome blisters for patch recordings using dehydration and rehydration as described previously^63^ with the following modifications. TMEM87A in buffer 1 was exchanged into soybean l-α-phosphatidylcholine (Soy PC, MillaporeSigma) with the addition of Biobeads SM2 (Bio-Rad) and an hour incubation at a protein:lipid ratio of 1:10 (corresponding to 0.4 mg purified TMEM87A and 4 mg of Soy PC lipid or 1:50 in buffer (5 mM HEPES pH 7.2, 200 mM KCl). TRAAK control proteoliposomes were prepared at 1:50 as described previously. Control liposomes were prepared from the same lipid and protocol with protein replaced with buffer 1.

### Electrophysiology

Proteoliposomes were thawed and dispensed in 0.5-1 μL drops on a 35 mm glass-bottom dish. The drops were dried in a vacuum chamber in the dark overnight. Proteoliposomes were rehydrated with 20 μL buffer (5 mM HEPES pH 7.2, 200 mM KCl). Each cake was firstly covered with a buffer drop and then let surface tension connect drops. Rehydrating proteoliposomes were placed within a humid chamber at 4 °C before patching. Recordings were made at room temperature using Clampex v.10.7 data acquisition software (as part of the pClamp v.10.7 suite) with an Axopatch 200B Patch Clamp amplifier and Digidata 1550B digitizer (Molecular Devices) at a bandwidth of 1 kHz and digitized at 500 kHz. A pressure clamp (ALA Scientific) was used to form seals. Pipette solution was 10 mM HEPES pH 7.2, 150 mM KCl, 3 mM MgCl_2_ and 5 mM EGTA. Bath solution was 10 mM HEPES pH 7.3, 135 mM NaCl, 15 mM KCl, 1 mM CaCl_2_, 3 mM MgCl_2_. Borosilicate glass pipettes were pulled and polished to a resistance of 2–5 MΩ when filled with pipette solution.

## Data availability

All data associated with this study will be publicly available. The TMEM87A model is in the Protein Data Bank (PDB) under 8CTJ, the final maps are in the Electron Microscopy Data Bank (EMDB) under EMD-26992, and the original micrograph movies and final particle stack are in the Electron Microscopy Public Image Archive (EMPIAR) under EMPIAR-XXXXX.

## Acknowledgements

The authors thank J. Remis, D. Toso, and P. Tobias for microscope and computational support at the Cal-Cryo facility of UC Berkeley. The authors thank all members of the Brohawn lab for helpful discussions and critical feedback on the project. S.G.B. is a New York Stem Cell Foundation-Robertson Neuroscience Investigator. This work was funded by the New York Stem Cell Foundation; NIGMS grant GM123496; a McKnight Foundation Scholar Award; a Sloan Research Fellowship; and a Winkler Family Scholar Award (to S.G.B.)

## Author contributions

L.Z. cloned, expressed, and purified proteins, prepared cryo-EM samples, and collected cryo-EM data. C.M.H. processed cryo-EM data and modeled the structure. L.Z. and C.M.H. performed electrophysiology. C.M.H., L.Z. and S.G.B analyzed data and wrote the manuscript. S.G.B. supervised the project.

## Declaration of Interests

The authors declare no competing interests.

---

**Supplementary Figure 1.**
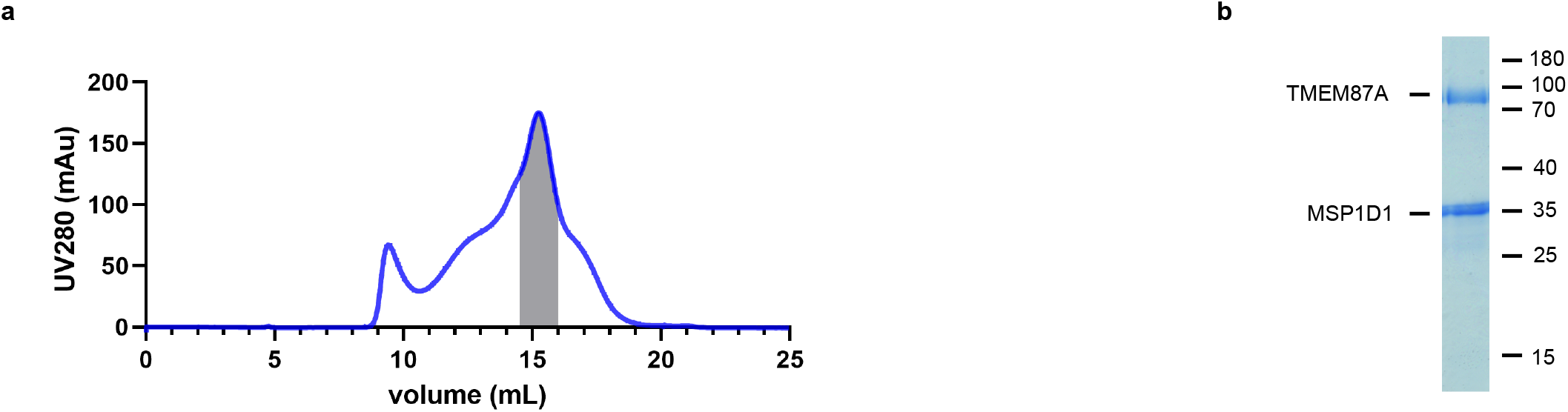
TMEM87A biochemistry. **(a)** Size-exclusion chromatogram and **(b)** Coomassie stained SDS-PAGE gel showing the purification and reconstitution of TMEM87A in lipid nanodiscs. Shaded region corresponds to pooled fractions collected. Samples were run on a Superose 6 column.

**Supplementary Figure 2.**
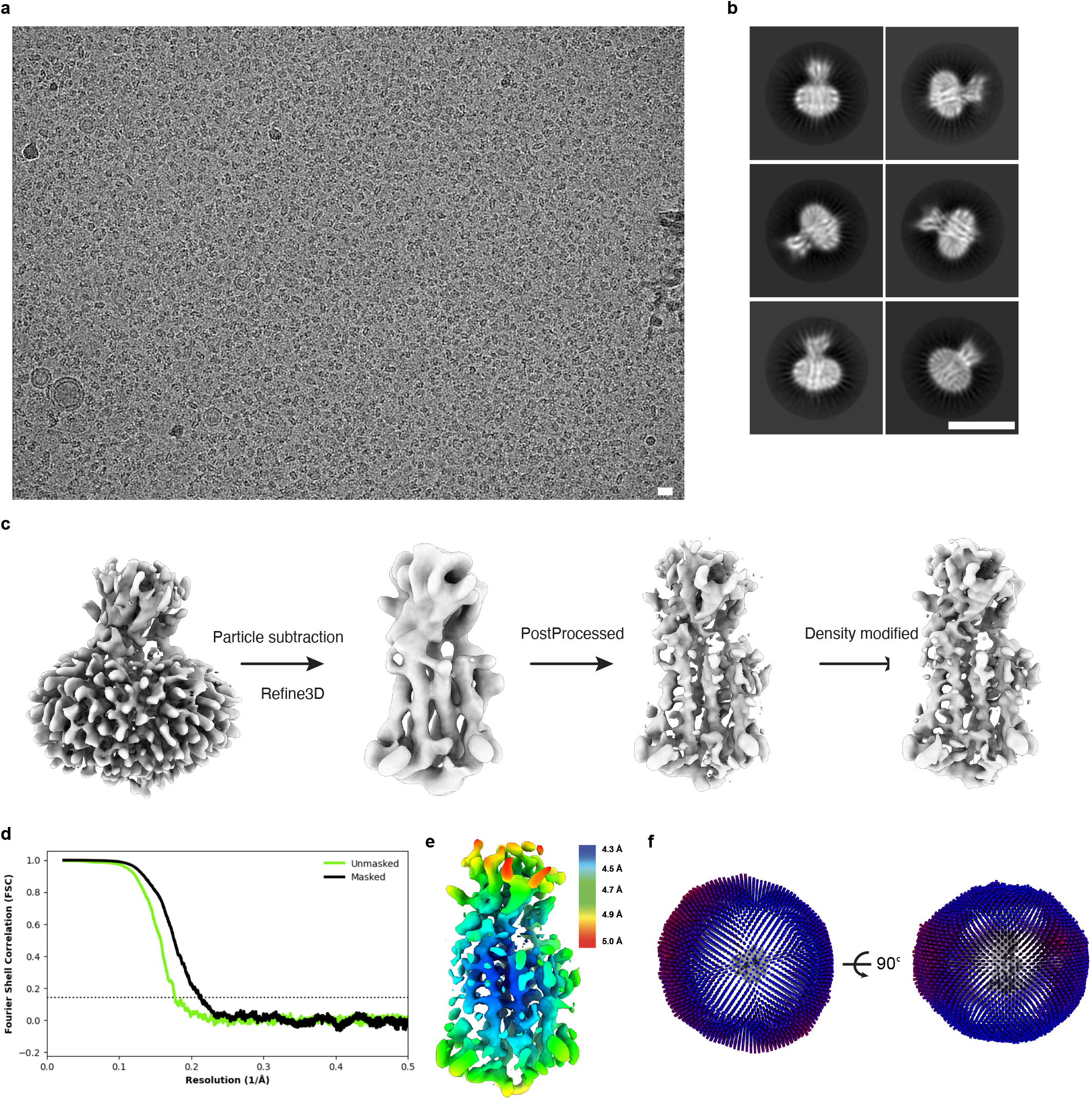
Cryo-EM data, processing, and validation. **(a)** Representative micrograph (scale bar=150 Å). **(b)** Selected 2D class averages (scale bar=100 Å). **c** Visual representation of final stages of the processing pipeline (see Methods for details). **(d)** Fourier shell correlations between the two corrected and masked (black) and unmasked (green) half maps. **(e)** Relion-estimated local resolution colored on the cryo-EM map in rainbow. **(f)** Angular distributions for particles used in the final refinement.

**Supplementary Figure 3.**
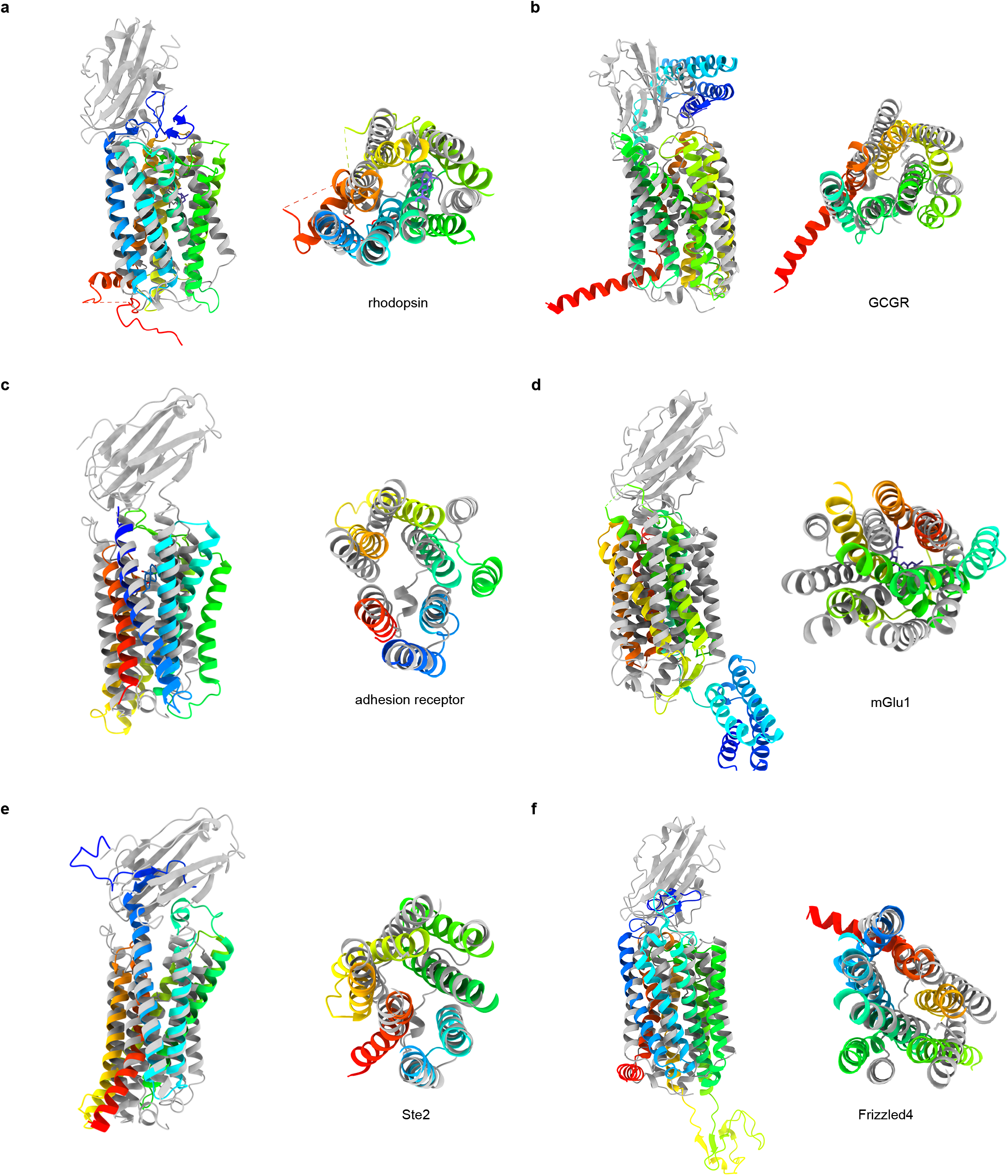
Comparison of TMEM87A and GPCR structures. **(a-f)** Overlay of TMEM87A (gray) with GPCR structures (rainbow from N-terminus (blue) to C-terminus (red)) viewed from the membrane plane (left) and above (right) in a 90° rotated view. **(a)** Class A GPCR bovine Rhodopsin (PDB 1F88) **(b)** Class B1 GPCR human glucagon receptor (GCGR) (PDB 4L6R) **(c)** Class B2 GPCR human adhesion receptor GPR97 (PDB 7D76) **(d)** Class C GPCR human metabotropic glutamate receptor 1 (mGluR1) (PDB 4OR2) **(e)** Class D GPCR yeast Ste2 (PDB 7AD3) **(f)** Class F GPCR human Frizzled 4 (PDB 6BD4).

**Supplementary Figure 4.**
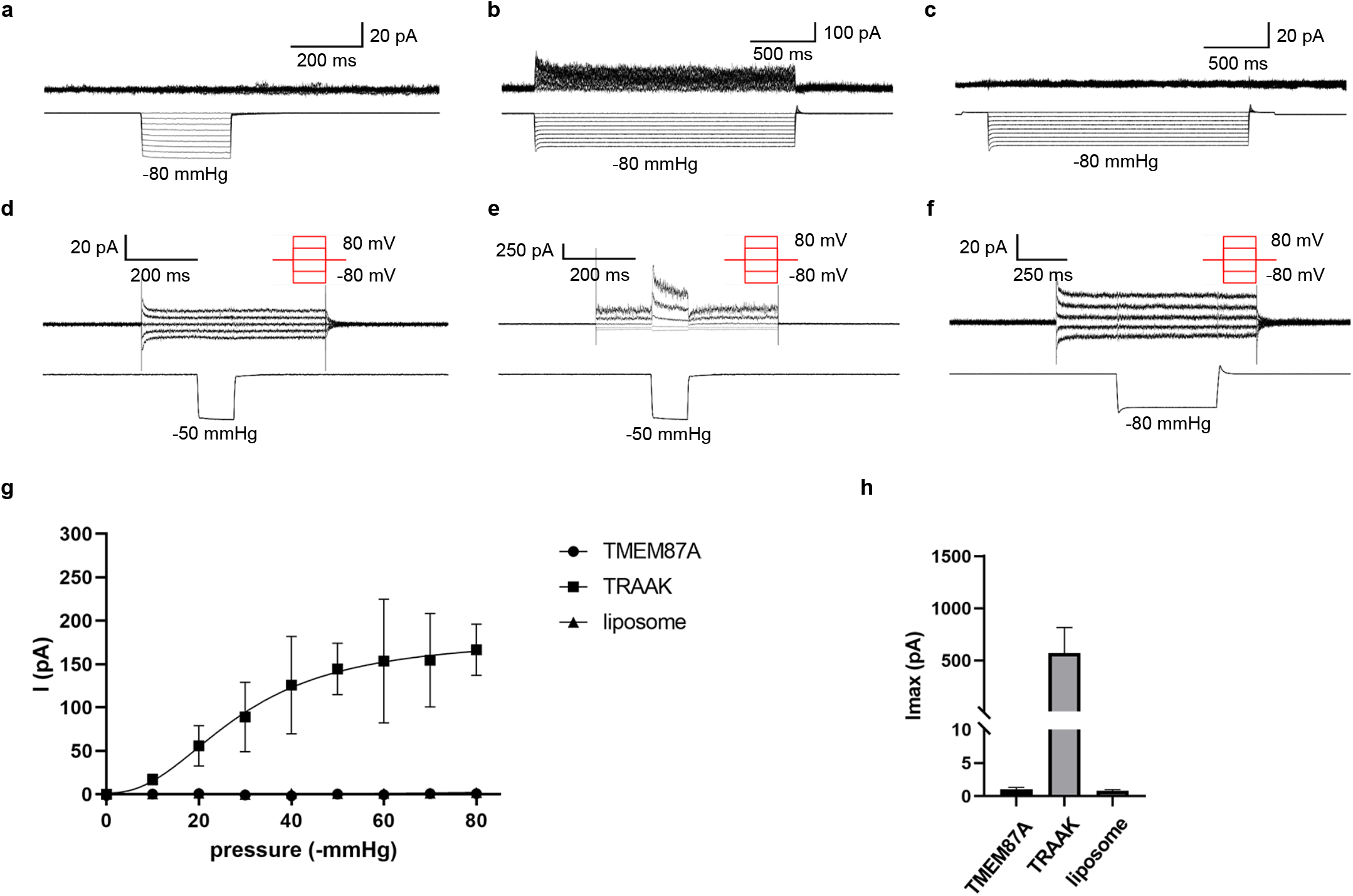
TMEM87A proteoliposome recordings. **(a-c)** Current (above) recorded during steps of increasing pressure (below) at a holding potential of 60 mV from a representative **(a)** TMEM87A-, **(b)** TRAAK-, and **(c)** mock-reconstituted proteoliposome patch. **(d-f)** Current (above) recorded during voltage steps (red schematic) during which a pressure step of −50 or −80 mmHg is applied from a representative **(d)** TMEM87A-, **(e)** TRAAK-, and **(f)** mock-reconstituted proteoliposome patch. **(g)** Average current vs. pressure plot from recordings as in **(a-c)** for TMEM87A (circles, n = 9), TRAAK (squares, n = 5), and mock-reconstituted proteoliposomes (triangles, n = 7). **(h)** Plot of maximum pressure (−80 mmHg) induced current observed from recordings as in **(a-c)** for TMEM87A (I_max_: 1.05 ± 0.52 pA, n = 4), TRAAK (I_max_: 572 ± 489 pA, n = 4), and mock-reconstituted proteoliposome patch (I_max_: 0.78 ± 0.60 pA, n = 7).

**Supplementary Figure 5.**
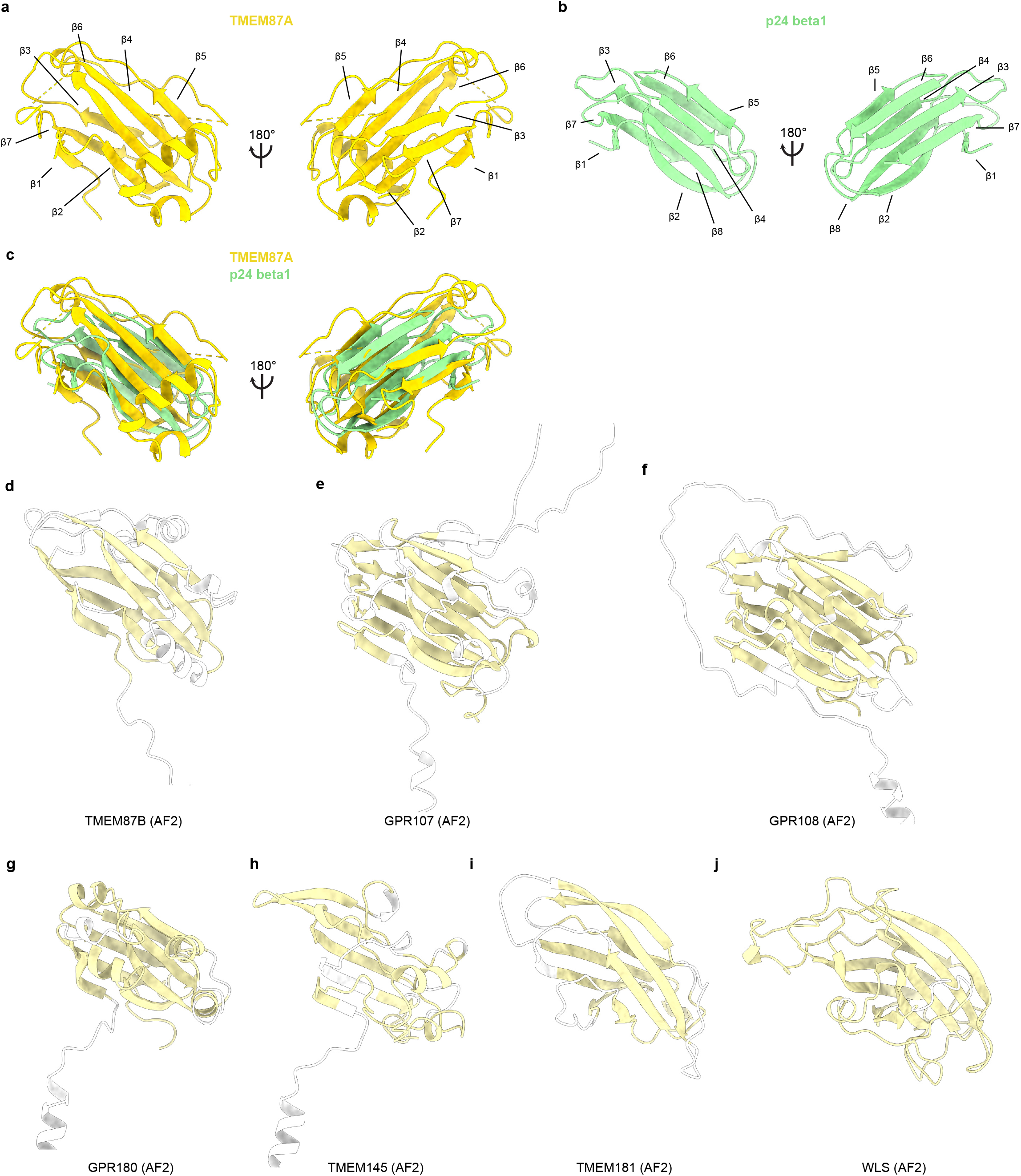
GOLD domain comparisons. **(a)** GOLD domain from experimental TMEM87A structure. **(b)** p24 beta1 GOLD domain (PDB 5AZW) colored light green. **(c)** Overlay of TMEM87A and p24 GOLD domains. **(d-j)** GOLD domains from AlphaFold2 predicted structures for TMEM87B, GPR107, GPR108, GPR180, TMEM145, GPR181, and WLS respectively, with GOLD domain colored light yellow and regions of low confidence (< 80 pLDDT) colored white.

**Supplementary Figure 6.**
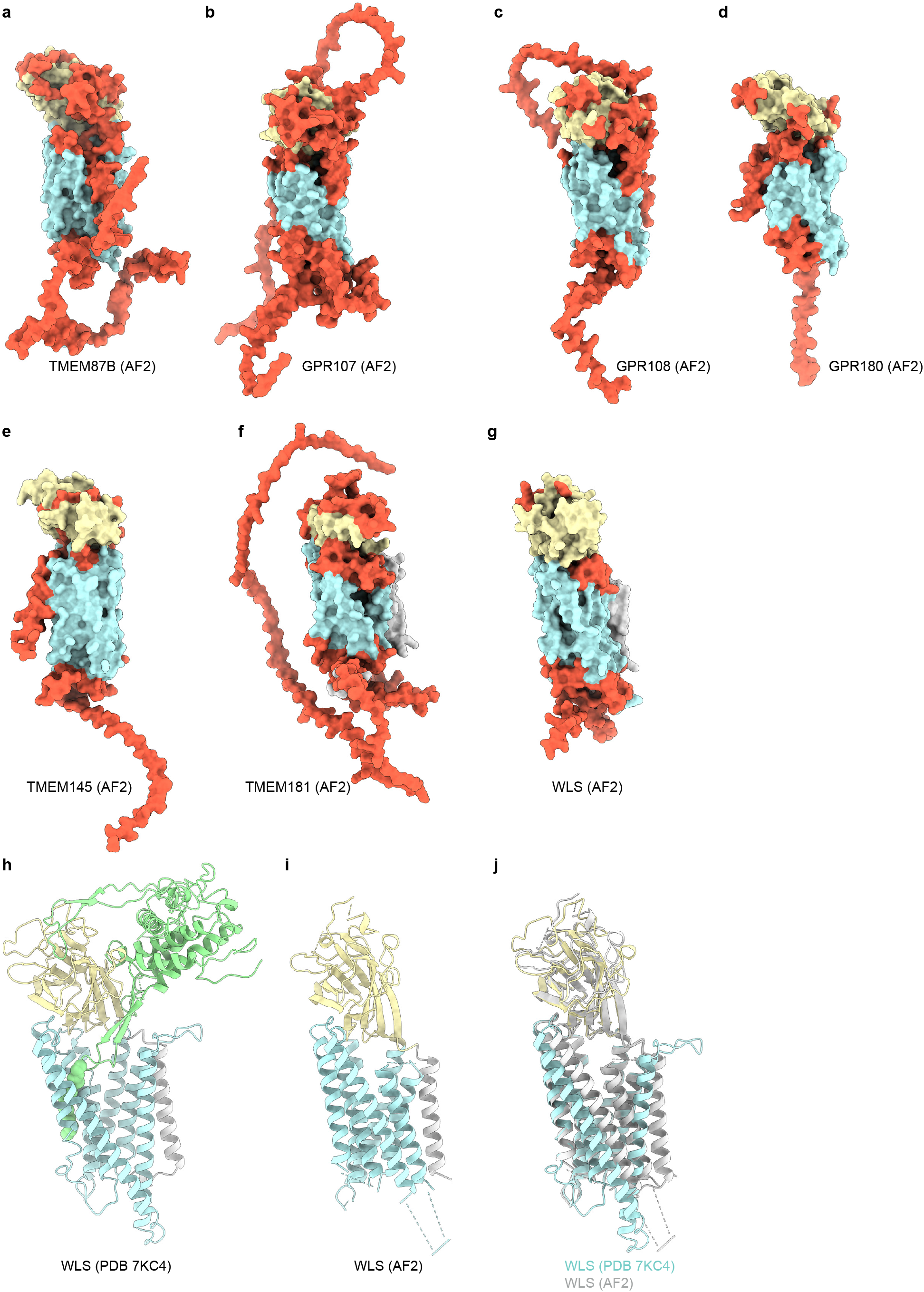
AlphaFold2 predictions of GOST family protein structures. **(a-g)** Surface representations of TMEM87B, GPR107, GPR108, GPR180, TMEM145, GPR181, and WLS AlphaFold2 predicted structures. GOLD domains are yellow, seven-transmembrane domains are blue, N terminal helices are gray (if present), and regions of low confidence (< 80 pLDDT) are orange. **(h)** Experimental WLS structure (7KC4) with GOLD domain yellow, transmembrane domain blue, N terminal helix gray, and Wnt8a green. **(i)** AlphaFold2 predicted WLS structure colored as **(h)**. Low confidence regions with pLDDT<80 are removed for clarity. **(j)** Overlay of experimental and predicted WLS structures.

**Supplementary Figure 7.**
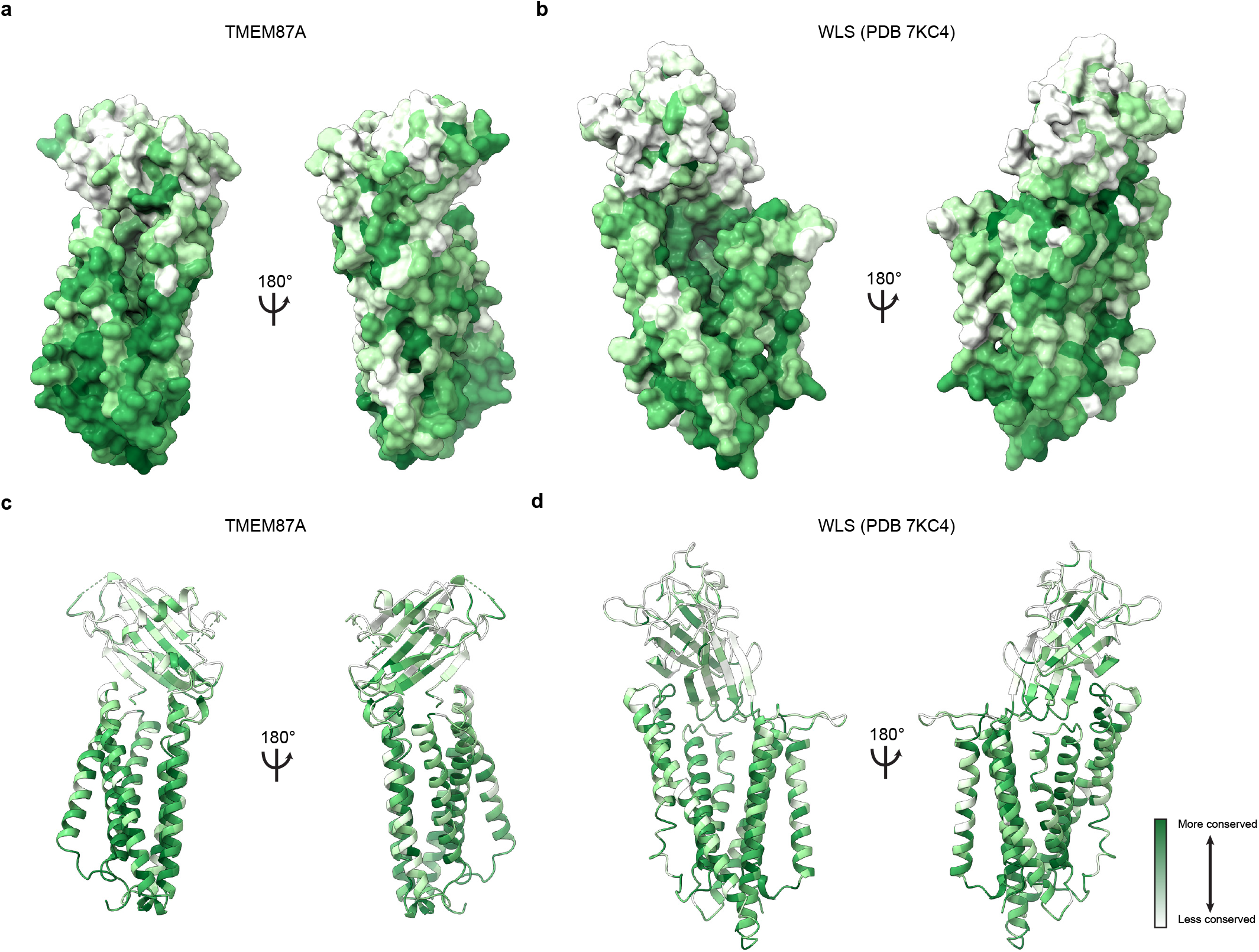
Conservation of TMEM87A and WLS. **(a,c)** TMEM87A structure colored according to conservation of each residue, from white (less conserved) to dark green (more conserved), on surface and ribbon representations respectively. **(b,d)** WLS structure colored according to conservation of each residue, from white (less conserved) to dark green (more conserved), on surface and ribbon representations respectively.

**Table S1.** Cryo-EM data collection, processing, refinement, and modeling data.

| Data collection | TMEM87A |
| --- | --- |
| PDB | 8CTJ |
| EMDB | 26992 |
| EMPIAR | XXXXX |
| Total movies # | 7060 |
| Selected movies # | 5898 |
| Magnification | 36,000 x |
| Voltage (KV) | 200 |
| Electron exposure (e/Å<sup>2</sup>) | 50 |
| Frame # | 50 |
| Defocus range (um) | -0.6 to -1.8 |
| Super resolution pixel size (Å<sup>2</sup>) | 0.5575 |
| Binned pixel size (Å<sup>2</sup>) | 1.115 |
| **Processing** |  |
| Initial particle images (no.) | 2,674,406 |
| Final particle images (no.) | 138,217 |
| Map resolution Masked (Å, FSC = 0.143) | 4.7 |
| Symmetry imposed | C1 |
| **Refinement** |  |
| Model resolution (Å, FSC = 0.143 / FSC = 0.5) | 3.9 / 5.9 |
| Map-sharpening B factor (Å<sup>2</sup>) | -421.5 |
| Composition |  |
| Number of atoms | 3312 |
| Number of protein residues | 401 |
| Number of ligands | 1 |
| R.m.s. deviations |  |
| Bond lengths (Å) | 0.004 |
| Bond angles (Å) | 0.855 |
| Validation |  |
| MolProbity score | 1.64 |
| Clashscore | 5.98 |
| Ramachandran plot |  |
| Favored (%) | 95.44 |
| Allowed (%) | 4.56 |
| Disallowed (%) | 0 |
| Rotamer outliers (%) | 0 |
| Mean B factor (Å<sup>2</sup>) |  |
| Protein | 153.9 |
| Ligand | 158.59 |

## Notes

### Competing Interest Statement

The authors have declared no competing interest.

